# A new pipeline to automatically segment and semi-automatically measure bone length on 3D models obtained by Computed Tomography

**DOI:** 10.1101/2020.06.06.137729

**Authors:** Santiago Beltran Diaz, Xinli Qu, Michael Doube, Chee Ho H’ng, John Tan Nguyen, Michael de Veer, Olga Panagiotopoulou, Alberto Rosello-Diez

## Abstract

The characterization of developmental phenotypes often relies on the accurate linear measurement of structures that are small and require laborious preparation. This is tedious and prone to errors, especially when repeated for the multiple replicates that are required for statistical analysis, or when multiple distinct structures have to be analysed. To address this issue, we have developed a pipeline for characterization of long-bone length and inter-vertebral distance using X-ray microtomography (XMT) scans. The pipeline involves semi-automated algorithms for automatic thresholding and fast interactive isolation and 3D-model generation of the main limb bones, using either the open-source ImageJ plugin BoneJ or the commercial Mimics Innovation Suite package. The tests showed the appropriate combination of scanning conditions and analysis parameters yields fast and comparable length results, highly correlated with the measurements obtained via *ex vivo* skeletal preparations. Moreover, since XMT is not destructive, the samples can be used afterwards for histology or other applications. Our new pipelines will help developmental biologists and evolutionary researchers to achieve fast, reproducible and non-destructive length measurement of bone samples from multiple animal species.

**Summary statement:** Beltran Diaz et al. present a semi-automated pipeline for fast and versatile characterization of bone length from micro-CT images of mouse developmental samples.

## INTRODUCTION

Skeletal measurement is the pillar of many research applications, such as developmental studies on limb patterning (Galloway et al., 2009; Summerbell, 1977) and growth (Marchini and Rolian, 2018; Rosello-Diez et al., 2017), main axis segmentation (Casaca et al., 2014; Wong et al., 2015), evolutionary studies (Kherdjemil et al., 2016; Sears et al., 2006; Sheth et al., 2012), disease modelling (Chen et al., 1999; Li et al., 1999; Rowe et al., 2018), adult phenotyping of mutant models (Boskey et al., 2003), etc. Whereas clinical musculoskeletal research often uses non-destructive imaging as routine (Cheng and Wang, 2018), fundamental evolutionary and development (evo-devo) studies often rely on differential staining of bone and cartilage (the so-called *ex vivo* skeletal preparations) (Mead, 2020; Rigueur and Lyons, 2014) and subsequent two-dimensional (2D) imaging for quantitative comparisons of the models of interest. Despite being broadly used, the skeletal preparation technique is ridden by several disadvantages. First, it is a destructive technique in the sense that the samples cannot be used for further histological or molecular applications. Second, it involves lengthy staining and clearing of cadavers, followed by laborious and damage-prone dissection of the skeletal elements of interest, in their preparation for imaging. Third, accurate measurements depend heavily on the imaged sample being positioned as flat as possible; otherwise the apparent length will be shorter than the real one due to parallax error. As a result, measurements are often prone to user error and require multiple measurements to calculate standard error. These limitations prompted us to seek alternative methods to measure bone length in a fast and reliable way, without destroying the sample.

X-ray microtomography (XMT) is a non-destructive imaging modality that uses radiographic projections taken at multiple angles to reconstruct two-dimensional tomograms (literally, slice images) whose pixel values represent the X-ray attenuation coefficient at each point in the imaged object (Elliott and Dover, 1982). Nowadays the process is run by computers, leading to the name of micro-computed tomography (μCT) It is common to arrange the tomograms into a stack to produce a three-dimensional (3D) image (Christiansen, 2016; du Plessis et al., 2017; Elliott and Dover, 1982). We reasoned that since XMT can be used to image undissected samples, it would allow us to scan multiple samples relatively fast, with the advantage of preserving their integrity in case they are needed for further processing. Moreover, computer-based image processing would in principle allow us to maximize the automation of the subsequent 3D reconstruction and measurements. Methods based on manual landmarking and measurement of the 3D models have already been developed, but we wanted to eliminate the human interaction component as much as possible. Our main goal, in summary, was to develop a pipeline to scan multiple whole-animal samples in a batch, and bulk-process the scans to extract linear measurements of the bones of interest. Minimal user intervention was the most important requirement, both to enable its use as a workhorse method in skeletal development labs, but also to eliminate any potential unconscious bias in the process. Within this general goal, we established three objectives: 1) to identify standard conditions (i.e. combination of scan resolution and analysis parameters) that yield low inter-batch variability; 2) to obtain a versatile pipeline that could be applied with minimal variation to a range of developmental stages; 3) to achieve enough precision to detect even small phenotypes, such as the 5-10% bone-length differences we have previously described with some of our models (Rosello-Diez et al., 2018; Rosello-Diez et al., 2017).

In XMT, the ability to independently analyse distinct tissues relies on their accurate separation through so-called segmentation (Bouxsein et al., 2010; Weissheimer et al., 2012). Since bone has a high mean atomic number and X-ray attenuation compared to other body tissues, it generates strong contrast in X-ray imaging modalities such as XMT and can be readily segmented through threshold-based methods where grayscale values determine what is bone tissue and what is background (Campbell and Sophocleous, 2014). There are several modalities of segmentation. Manual segmentation involves the manual selection of the areas of interest section by section, and is therefore quite laborious and subjective, thus prone to user error (Rathnayaka et al., 2011). Semi-automated methods, on the other hand, use algorithms like edge detection (Rathnayaka et al., 2011) and/or local differences in grey values (Zhang et al., 2010) with some user input for initial parameters. Another common method is automated segmentation, whereby image-processing algorithms are used to segment elements of interest with minimal to no-user interaction (Heidrich et al., 2013; Okada et al., 2008; Šajn et al., 2007; Yiannakas et al., 2016). Algorithm-based automatic segmentation, however, requires the user to have programming knowledge and a thorough understanding of mathematical algorithms related to the image processing software being used (Rathnayaka et al., 2011).

There are a wide range of software solutions that can analyse XMT data in the form of digital imaging communications in medicine (DICOM) files to segment a variety of high-contrast tissues like lungs (Reynisson et al., 2015; Weissheimer et al., 2012), liver (Huhdanpaa et al., 2011; Okada et al., 2008) and bone (Mehadji et al., 2019; Rios et al., 2014; Taghizadeh et al., 2019). After some pilot testing of both open-source and commercial solutions, we settled on the Mimics Innovation Suite (Materialise, Leuven, Belgium) as the one that most readily suited our needs. Mimics has been previously benchmarked against other programs like Syngo (An et al., 2017), OsiriX (Reynisson et al., 2015) and ITK-snap (Weissheimer et al., 2012), and some of its key features are its flexibility, ease of use, sensitive and controlled segmentations (Reynisson et al., 2015; Weissheimer et al., 2012) and the possibility to integrate Python scripting modules to further extend its automation capabilities.

Here we present a semi-automated analysis pipeline for the fast and robust characterisation of long-bone length, using two solutions that can be adopted by non-experts: 1) Python scripting and segmentation tools of the commercially available software package Mimics; 2) a standardised pipeline in the ImageJ plugin BoneJ. We report the advantages and caveats of each method.

## RESULTS

### A script for bone-length measurement on XMT scans with minimal user input using Mimics

After testing multiple software solutions, we chose a commercial one (Materialise Mimics Research software) to develop the initial pipeline, based on the promising initial results and the possibility of automation via scripting. We thus developed a Python script that utilises Mimics capabilities to segment and measure the mouse bones of interest (humerus, radius, ulna, tibia and sometimes clavicle) from CT scans. This script is called BASILISC (<u>B</u>one <u>A</u>utomated <u>S</u>egmentation and <u>I</u>nteractive <u>L</u>ength <u>I</u>nterrogation on Standardized computerized scans). BASILISC is available in Github (www.github/rosellodiez/Basilisc), and designed to run in the Materialise Mimics Research software v.18 to 21, and hence there are attributes that are specific to this program. The script can be divided into 4 main sections: thresholding, landmarking, 3D modelling and measurement & export (Fig. 1A). See Supplementary Video 1 for an overview of the whole procedure.

**Figure 1.**
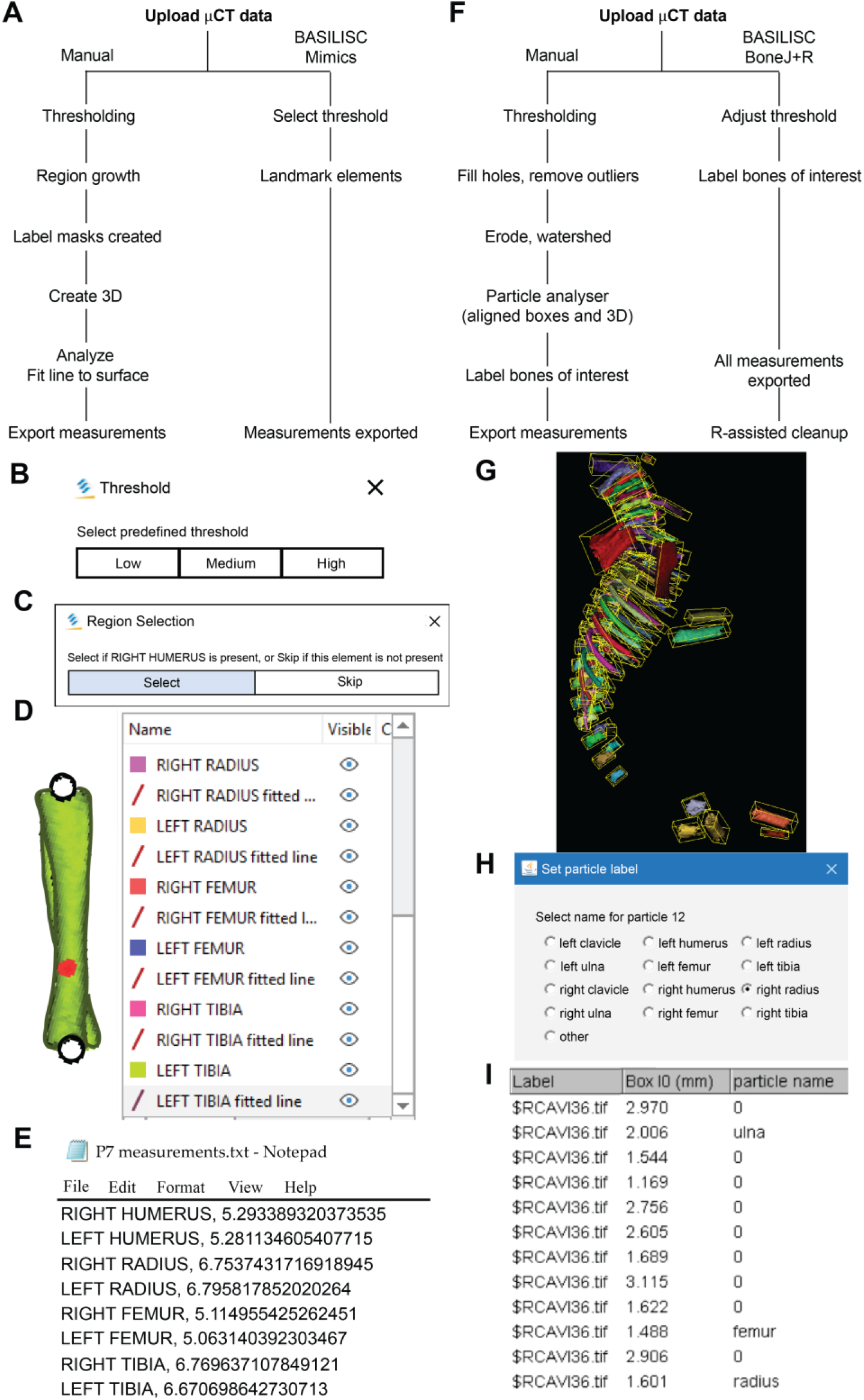
Bone Automated Segmentation and Interactive Length Interrogation of Standardized computerized scans (BASILISC). A-E) Mimics procedure. A) Diagram depicting the Mimics procedure followed by the script. B-E) Representative screenshots of key steps in the process: threshold preselection and segmentation (B), element seeding (C), fit-to-surface line fitting (D), table export (E). F-I) BoneJ procedure. F) Diagram depicting the BoneJ procedure followed by the macro. G) Typical output of the *Particle Analyser* module, as visualized with ImageJ’s 3D Viewer. Aligned boxes appear in yellow. H-I) The new BoneJ module *Label Elements (3D)* allows the user to interact with the 3D viewer to select the bones of interest, select or write a label for them (H) and append that label to the *Results* table (I). Note: the outlines and fonts of the screenshots have been redrawn for aesthetic and readability purposes.

The first Python command in BASILISC segments all the skeletal elements, using a global threshold for bone tissue (Fig. 1B). To increase its applicability to different developmental stages and scanning conditions, BASILISC was designed in such a way that the user can select among three pre-defined thresholds (Low, Medium, High, ranging from ~200 to 500 Hounsfield Units (HU)) via a pop-up menu. These pre-defined values can be easily changed within the script (see Methods). The next section of the script is landmarking, which uses Mimics tools to segment and uniquely label all bones of interest via a pop-up menu (Fig. 1C). The user is prompted to select a region (landmark) of the indicated element by simply clicking on it on one of the 2D views of the sample. BASILISC will automatically label and segment the selected element without further user interaction. In the third section of the script, once all bones of interest have been segmented and labelled accordingly, a 3D model of each skeletal element is created (Fig. 1D). Then BASILISC automatically fits a ‘line to surface’ running from end to end along the centre of each element. In the last section, the script automatically obtains the length of the fitted lines and saves the measurements to a comma-separated-values (csv) text file (Fig. 1E).

### A pipeline for streamlined bone length measurement in the open-source ImageJ plugin BoneJ

We also developed a similar pipeline to semi-automatically measure bone length using open-source software, as a broadly useful approach. We chose ImageJ (https://imagej.nih.gov/ij/), a popular Java-based image analysis program developed at the U.S.A. National Institutes of Health, as a platform for such a solution. Importantly, one of us (M.D.) previously developed an ImageJ plugin (BoneJ) with modules designed for analysis of bone geometry (Domander et al., 2021). Its ability to load and segment micro-CT data was the starting point we required. We also developed new modules and new capabilities for existing ones to suit our purposes (see Material and Methods). The pipeline can be divided into loading, segmentation, fitting a minimal bounding box to each element and measurement of said box (Fig. 1F). A brief overview follows.

We first loaded the whole data set (DICOM format) into FIJI (a package that includes ImageJ and multiple biology-oriented plugins for image analysis (Schindelin et al., 2012)). We then proceeded with segmentation (i.e. the isolation of skeletal elements form everything else). We chose to use a maximum entropy method because in our experience it offers the best robustness, acceptable performance with low-contrast images, and good noise tolerance (Kapur et al., 1985). The macro prompts the user to enter a value for the lower threshold, which needed to be adjusted depending on the stage being analysed, and for our scans it ranged between 280 and 500 HU. We then used the *Fill holes* tool (2D version) to close solid structures such as the marrow, which facilitates downstream analysis. To clean up the data, we used the *Remove outliers* tool to remove small radio-dense objects that may appear in some images, followed by the *Erode* and the *Watershed* commands (2D versions), to facilitate the separation of structures that are frequently close to each other (e.g. radius and ulna). Finally, to obtain the length of the different bones without the time investment of selecting points manually on a 3D object, we first attempted to use the *Particle Analyser* tool of BoneJ (Doube, 2021) to calculate approximations of bone dimensions and plot them on 3D models of the segmented elements (Suppl. Fig. 1A). Suppl. Fig. 1B shows examples of the common measurements and fitted shapes that could be computed with BoneJ before this project was started, namely the maximum Feret’s diameter, a fitted ellipsoid and the three moments of inertia. The Feret’s diameter (also known as the maximum calliper diameter) is the longest distance between any two points on the particle’s surface. For rod-like structures with relatively flat ends, Feret’s diameter is a reasonable approximation of length. However, we noticed that in the case of real-life bones, the Feret’s diameter tends to find a diagonal that is not aligned with the main bone axis, leading to an overestimation of the length (Suppl. Fig. 1B-B’’, green arrowheads). Additionally, it is implemented as a brute-force method so it is computationally demanding. The fitted ellipsoid is the best-fit ellipsoid to the particle’s surface mesh. Since long bones are not ellipsoidal in shape, the ellipsoid fit is not precise and the fitted ellipsoid’s long axis tends to be longer than the bone’s (Suppl. Fig. 1B, blue mesh). Finally, the moments of inertia are defined with respect to the three mean principal axes of the segmented element (longest, shortest and middle, Suppl. Fig. 1B, red lines). Therefore, we reasoned that the intersections between the longest axis and the surface of the 3D model of the skeletal element could be used to define a straight line approximating bone length. However, we noticed that while the principal axis ran reasonably well along the centre of bones such as the femur (e.g. Suppl. Fig. 1B), this was not the case for other bones such as the humerus, where the identified principal axis formed a steep angle with the line that a human user would use to measure bone length (Suppl. Fig. 1B’’). In summary, BoneJ was able to automatically provide good approximations of bone length for only some bones, depending on their shape.

Given these limitations, we developed a new capability for BoneJ’s *Particle Analyser*, to generate and measure *Aligned boxes* fitting each element (Fig. 1G). The new tool uses the bone’s inertia tensor to define the three principal axes of each element, and then generates the minimum-size cuboidal box that fits the skeletal element and that is aligned with the principal axes. The coordinates and dimensions of these boxes are then appended to the *Results* window in ImageJ. One challenge of this approach is that for whole-body scans, hundreds of elements are segmented and measured (Fig. 1G), and the numeric IDs assigned to the different bones are not the same from sample to sample, precluding the streamlined export of only the measurements of interest. While filtering by volume in the *Particle Analyser* tool allows reducing the number of elements to certain extent, it is often impossible to reduce it only to the bones of interest. Therefore, we generated a new analysis module for BoneJ, *Label Elements* (*3D*), which allows the user to interact with the 3D Viewer in order to Control-click the elements of interest, subsequently selecting or writing the label for the clicked element in an iterative manner (Fig. 1H). That label is appended to the Results window (Fig. 1I), which is subsequently exported as a csv file. Lastly, we developed an R-script to clean up the data tables (i.e. to remove all the rows but the ones of interest), sort the remaining values in alphabetical order and finally to bind several data tables together, if required (see Methods).

### Standardized conditions to achieve robustness to batch effect at multiple stages

A semi-automatic protocol to measure bone length would only be useful if it yielded consistent measurements for a given sample, scanned and analysed repeatedly on different days. We thus explored different scan resolutions (20 and 40 μm) to analyse at least three technical replicates per resolution, obtained from two different postnatal day (P) 7 mouse specimens (see Methods), and assessed the reproducibility of the results (Fig. 2). We report the Mimics and BoneJ results separately below.

**Figure 2.**
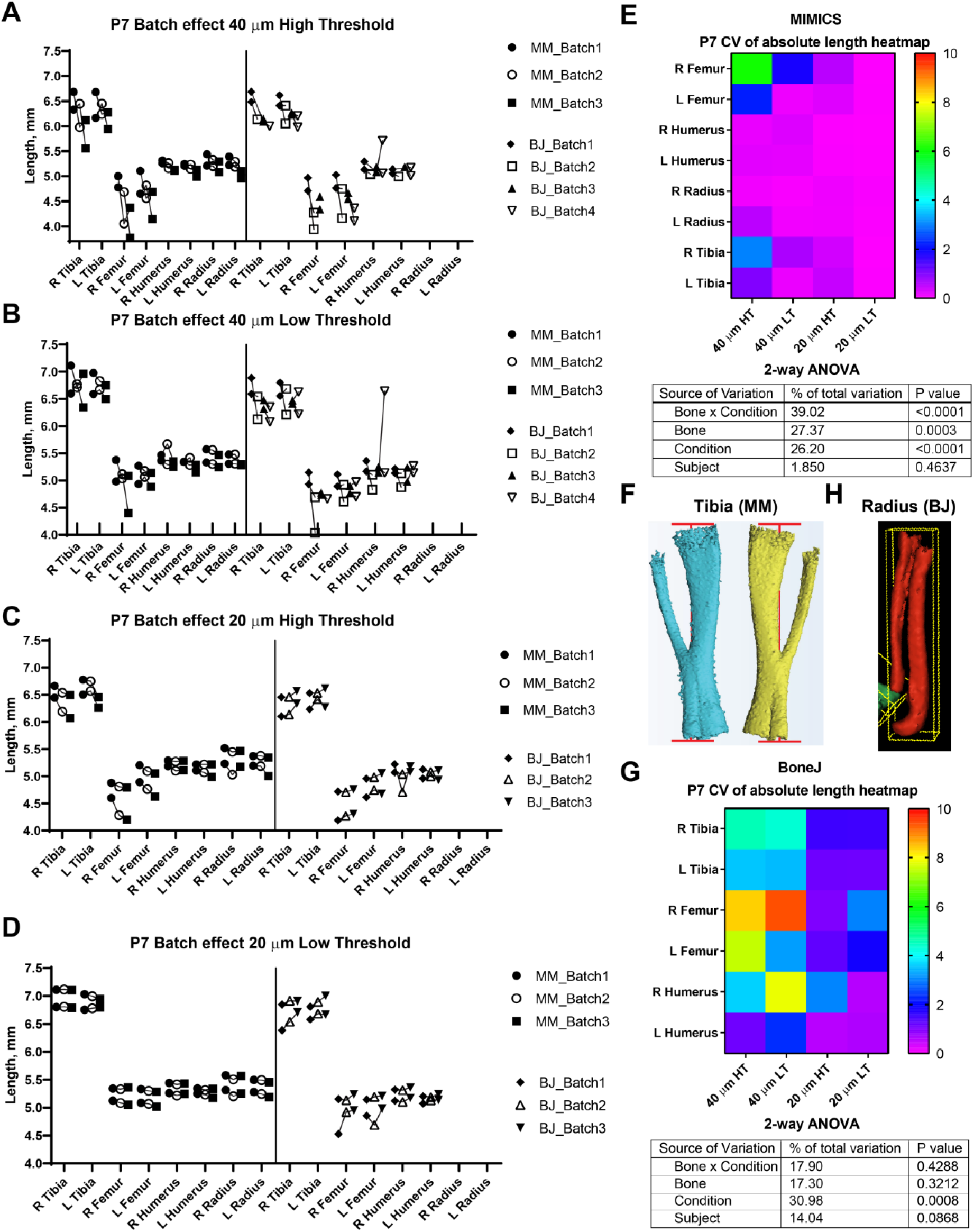
Assessment of batch effect for multiple P7 bones across different scan and analysis conditions. A-D) Measured length for the indicated bones of two P7 mouse pups, each scanned on three different days (triplicates joined by lines) at either 40 (A, B) or 20-μm resolution (C, D), and analysed with either a high (A, C) or a low (B, D) minimum threshold. L, R: left, right. E and G) Top: Heatmap for the Coefficient of Variability (CV, %) between the three batches of the indicated measurements, for the Mimics (E) and the BoneJ (G) pipelines. Bottom: 2-way ANOVA table showing the contribution and associated p-value of each source of variation of the experiment. F, H) Representative examples of the generated 3D models (left and right from the same specimen) and their fitted lines/boxes, for tibia using Mimics (F) and for radius/ulna using BoneJ (H).

In the Mimics pipeline, 40-μm scans showed relatively high inter-batch effect, especially for hindlimb bones, regardless of the threshold (Fig. 2A-B, left), whereas 20-μm scans yielded more consistent measurements, including hindlimb bones, especially for the lower threshold (Fig. 2C-D, left). To compare the batch effect more quantitatively, we then calculated the coefficient of variability (CV) for each bone’s measurements across the three batches, and compared the CV for the different conditions and bones. A 2-way analysis of variance (ANOVA) showed that there was a significant effect of the imaging & analysis conditions, although the extent of it was likely distinct for the different bones (Fig. 2E). In summary, these results identified a 20-μm resolution and a 398-HU threshold as the optimal conditions to minimize inter-batch variability in this type of samples (i.e. P7 mouse long bones). Importantly, this protocol succeeded to separate radius from ulna in most cases, despite their proximity, with only a few scans requiring the manual use of *Split Mask*. However, it was not able to separate tibia from fibula (Fig. 2F), potentially affecting tibial length measurement (see Discussion below).

With BoneJ, the 20-μm resolution also led to lower inter-batch variability, but in this case there was no major difference between the high and low threshold for any resolution, although the high threshold showed a trend towards less variability (Fig. 2A-D, G). As in the Mimics case, the femur was the long bone most affected by high variability, especially under suboptimal conditions of resolution (Fig. 2G). One obvious difference with the Mimics pipeline is that, in the BoneJ pipeline, radius and ulna were segmented together in most cases (Fig. 2H). In the absence of a *Split mask* function that could be applied to separate those elements in a quick an efficient way, we concluded that radius measurements could not be included in the analysis.

In order to test the versatility of BASILISC across developmental stages, we performed a similar battery of scan & measurement analyses exploring different scan resolutions and threshold values at embryonic day (E) 17.5 (Fig. 3). With Mimics, most of the conditions performed similarly in terms of reproducibility across batches, except for low resolution and threshold, for which some femora were not properly segmented and as a consequence their length was overestimated (Fig. 3A-D). Although there was no overall difference in the CV across conditions (Fig. 3E), the data trends suggested that a 398-HU threshold outperformed a 226-HU threshold and length variation due to differences in scan resolution were minimised with a 398-HU threshold. Importantly, tibia and fibula were again segmented together no matter the conditions of analysis (Fig. 3F).

**Figure 3.**
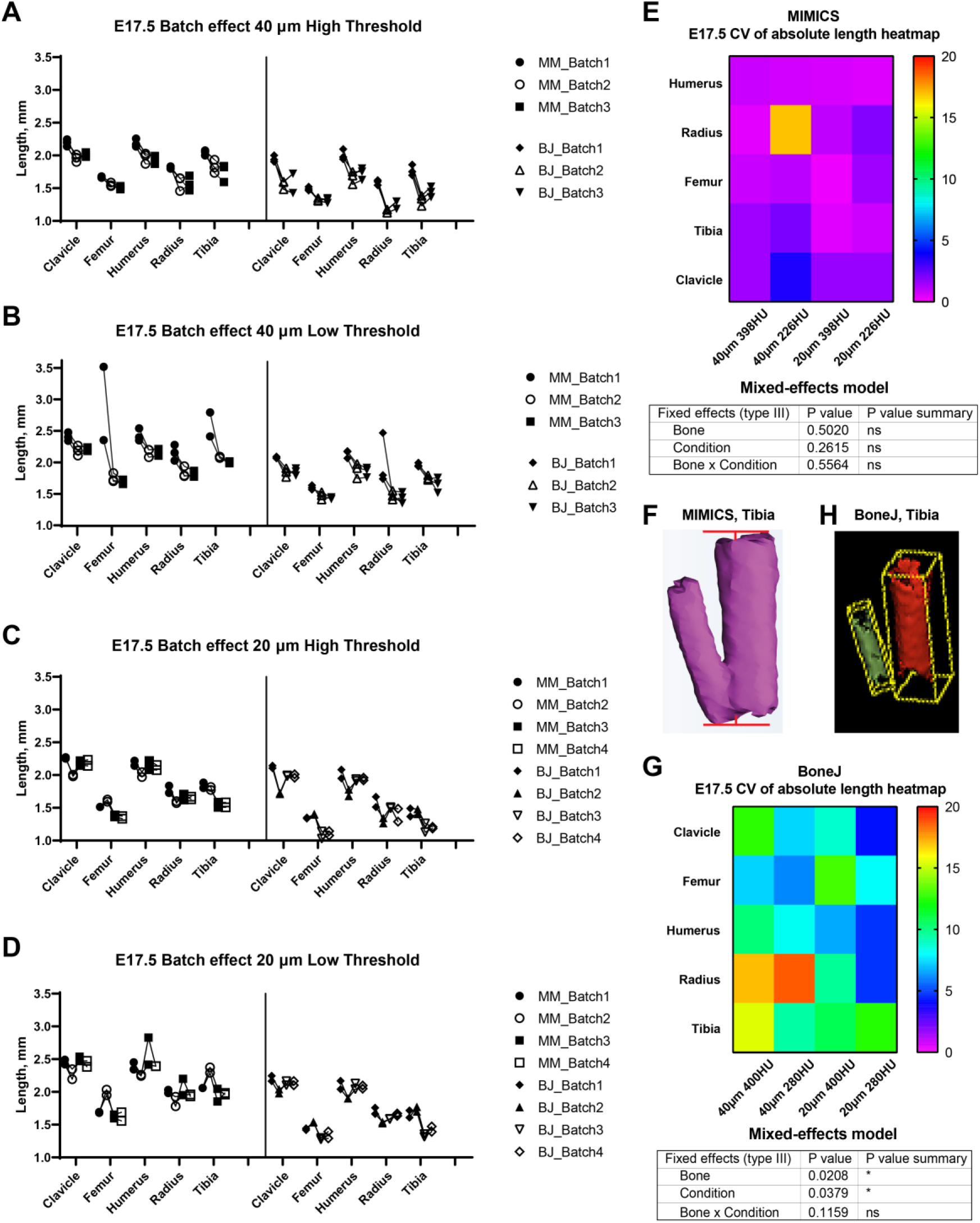
Assessment of batch effect for multiple E17.5 bones across different scan and analysis conditions. A-D) Measured length for the indicated bones of two or three E17.5 mouse foetuses, each scanned in triplicate (or quadruplicate) at either 40 (A, B) or 20-μm resolution (C, D), and analysed with either a high (A, C) or a low threshold (B, D). E, G) Top: Heatmap for the Coefficient of Variability (CV, %) between the batches of the indicated measurements for the Mimics (E) and BoneJ (G) pipelines. Bottom: mixed-effects model table showing the contribution and associated p-value of each source of variation of the experiment. F, H) Representative example of the generated tibial 3D models and its fitted line (F) or aligned box (H).

With the BoneJ pipeline, variability was somewhat higher, although again high resolution and low threshold tended to yield more robust results (Fig. 3A-D, G). At this stage, radius and ulna were, as with Mimics, individually segmented in all cases. One important difference with the Mimics pipeline is that tibia and fibula were not always segmented together. Namely, low resolution and high threshold led to separated tibia and fibula in only ~half of the cases and therefore to high variability (Fig. 3A, G); low resolution and low threshold led to fibula and tibia always segmented together, such that there was less variability in the measurements, although the measurement itself was somewhat overestimated (Fig. 3B, G); high resolution and high threshold led to separated tibia and fibula in ~88% of the cases but also to highly eroded skeletal elements (Fig. 3H), which was associated with intermediate variability (Fig. 3C, G); and high resolution and low threshold also led to separated tibia and fibula in ~88% of the cases but with less eroded surfaces, which likely corresponds to more exact measurements (Fig. 3D, G).

### Internal consistency across stages, scan resolutions and segmentation thresholds

Besides inter-batch variability, a good analysis pipeline should yield low intra-scan variability. In other words, the ratios between two specific bones should be very similar across different scans. One of the advantages of working with paired bones is the possibility of assessing internal consistency of the BASILISC method by measuring the left/right ratio for each bone and condition. We therefore calculated a left/right ratio for the P7 samples, including replicates, to determine how close the ratio was to the hypothetical value of 1 (i.e. equally long left and right paired bones) and how much variability there was between replicates. As shown in Fig. 4A-B, for P7 samples 20-μm scan resolution low threshold again had the lowest inter-batch variability and the L/R ratio was remarkably close to 1. As parameters moved away from these optimal settings, there were several bones (typically femur, tibia and sometimes radius) for which either the average value was not as close to 1 as for other bones, and/or the variability between batches was higher than 5% (Fig. 4A-B and S2). Similarly, we calculated internal ratios for E17.5 bones to determine optimal scan and segmentation parameters. In this case we chose intra-limb ratios (humerus/radius and femur/tibia) as a normalisation approach that could be achievable in the case that contralateral bones were not available (as it was our case for these scavenged samples). With this approach, the parameter of interest to estimate the precision of the approach was the variability of each measurement across replicates. High resolution and low threshold minimized variability, and in all conditions the Mimics pipeline was more accurate (Fig. 4C-D, S2).

**Figure 4.**
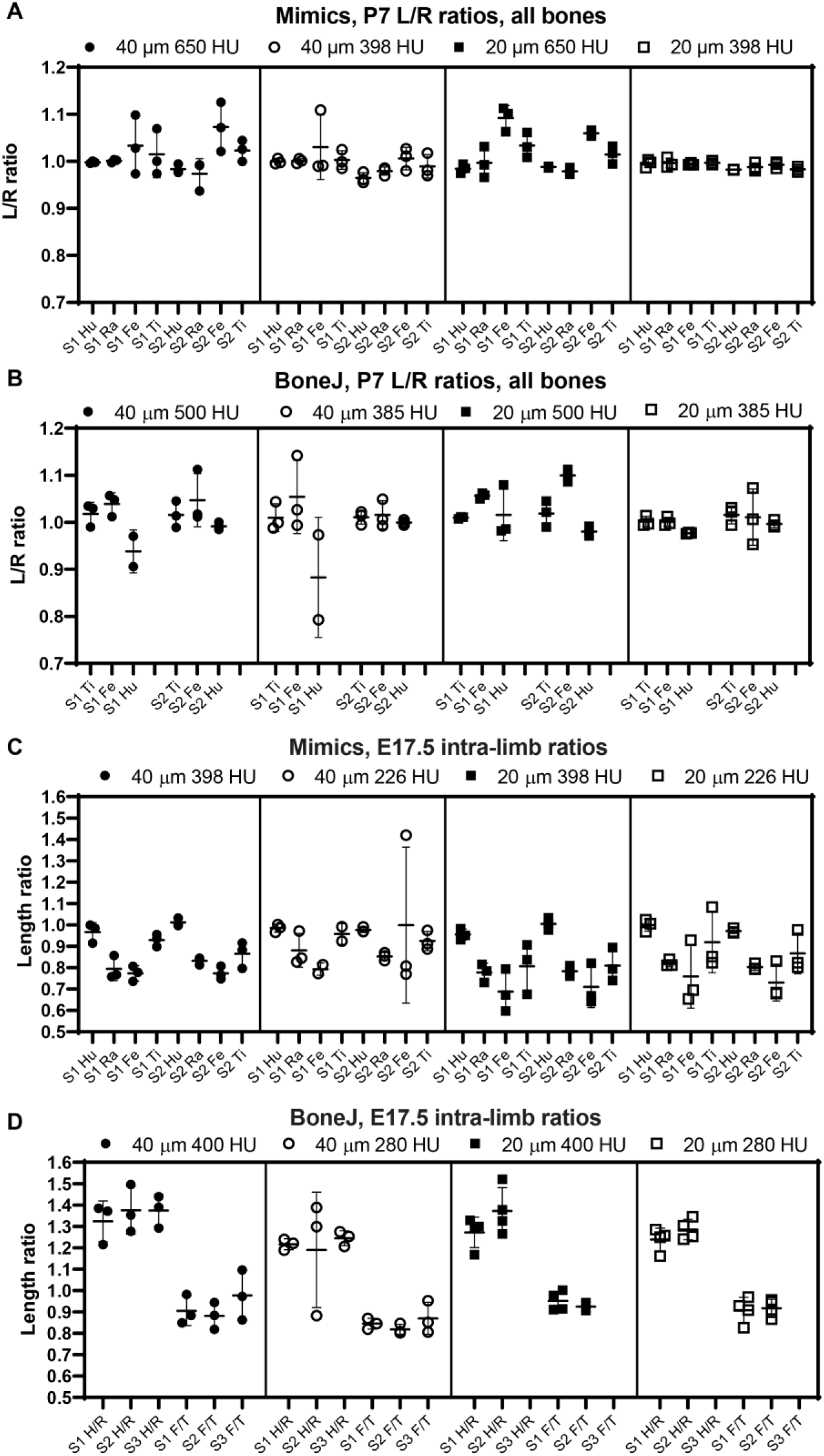
The comparison of intra-specimen ratios reveals the most reproducible conditions for scan and analysis. A-B) Left/right ratio of bone length (mean±SD) for the indicated bones and conditions at P7, using Mimics (A) or BoneJ (B). Hu/Ra/Fe/Ti, Humerus/Radius/Femur/Tibia. C-D) Similar to A-B), except that the ratios shown are Humerus/Radius (H/R) and Femur/Tibia (F/T). S1-S3, specimens 1 to 3.

In summary, these results suggest that, as expected, 20-μm resolution is preferred to 40-μm in order to achieve more robust measurements. In terms of threshold, a higher threshold is preferred to segment out bones that are close to each other, but if the separation is not always achieved this leads to higher variability of the measurement. We thus concluded that the exact threshold needs to be determined for each scanner and/or scanning condition. Of note, the threshold can be easily changed in the Mimics script and the ImageJ macro that we present here.

### Correlation bone lengths obtained via Mimics, BoneJ and skeletal preparations on the same samples

We next compared the bone lengths obtained by BASILISC (Mimics pipeline) with the lengths obtained from the same samples via skeletal preparations and digital measurement of photographed bones, a method frequently-used in developmental biology studies. We used eight long bones from three different specimens at P7. The linear relationship between both measurements was very good in all conditions (Fig. 5A, p-value for Pearson correlation <0.0001 in all cases), and the slopes were not significantly different (p=0.3298), with an average common value of 0.9642. As expected, however, the BASILISC measurements that used lower thresholds tended to overestimate bone length (as the resulting 3D model includes less dense tissue), as indicated by the differences in the intercepts with the axes (Fig. 5A, p<0.0001). Overall, the conditions that yielded measurements with better correlation to the skeletal preparations were 20-μm resolution and low threshold. We thus restricted our next analysis to these conditions, comparing BoneJ and Mimics BASILISC with the skeletal preparations (Fig. 5B). Both correlations showed very similar slopes and Y-intercepts with the P7 samples. Both BASILISC approaches led to greater measurements than the skeletal preparation, especially in the case of Mimics (Fig. 5B).

**Figure 5.**
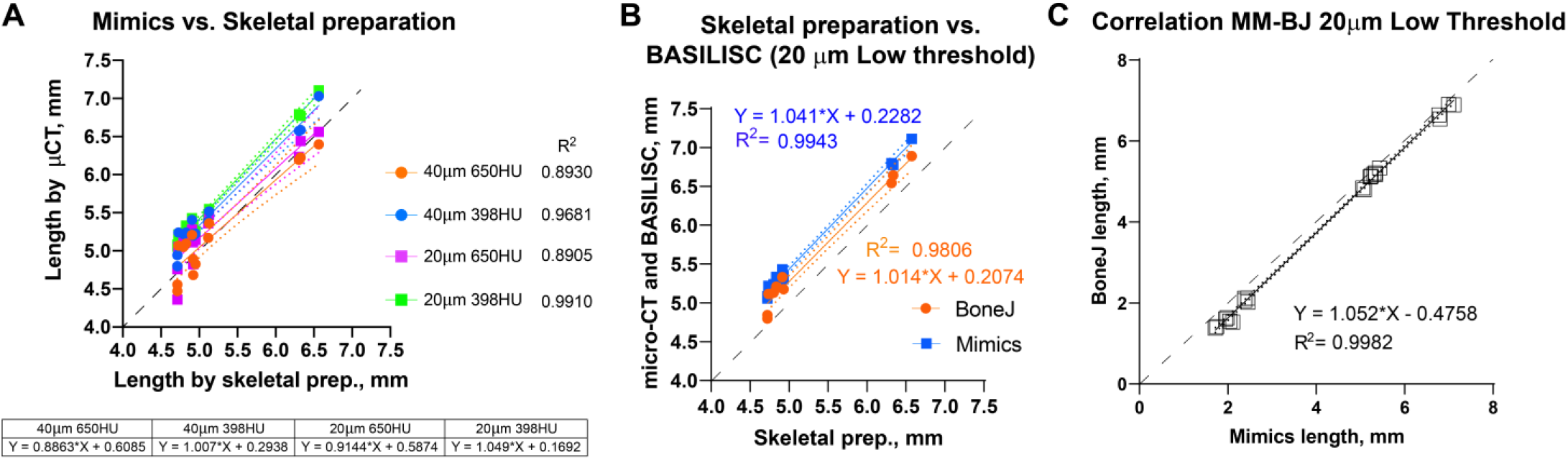
Correlation between BASILISC methods and with measurements obtained via skeletal preparations. A) Bone length measurements obtained via skeletal preparations (prep.) (X axis) and Mimics BASILISC (Y axis), for different combinations of imaging resolution and segmentation threshold (averages of 2-3 technical replicates are shown). Solid lines represent the regression line for each combination. The coefficient of determination (R^2^) is indicated. The table shows the slope and Y-intercept for each of the conditions used. B) Bone length measurements obtained via skeletal preparations (prep.) (X axis) and via micro-CT with Mimics and BoneJ versions of BASILISC (Y axis), obtained at 20-um resolution and low threshold. C) Correlation between bone length measurements of 22 bones (10 from E17.5 embryos and 12 from P7) obtained with our Mimics (X axis) and BoneJ (Y) pipelines. Each Mimics-BoneJ comparison was done on the same scan batch. In A-C), the dashed black line represents a 1:1 correlation as a reference, and the dotted ones delimit the 95% confidence interval of the regression.

As the last test for our pipelines, we compared the measurements obtained by Mimics and BoneJ from the same samples and scan batches, covering a broader range of stages, i.e. from E17.5 to P7. We focused on the conditions that yield most robust results, namely 20-μm resolution and low threshold. As seen in Fig. 5C, there is remarkable correlation between both methods (R^2^=0.9982), although BoneJ leads to smaller lengths than Mimics (which itself leads to greater lengths than the skeletal preps, Fig. 5A, B).

We concluded that both pipelines were suitable for high-throughput analysis of bone length from whole-body scans, with minimal user intervention. Next, we tested these methods in proof-of-principle studies.

### Application of BASILISC to the analysis of genetic mouse models with altered skeletogenesis

To test the utility of the BASILISC pipeline for the detection of skeletal phenotypes that affect bone length, we applied it to one of our models of limb asymmetry that we recently reported (Ahmadzadeh et al., 2020). In this model, diphtheria toxin expression is activated in an inducible and reversible manner in the cartilage template that drives growth of the left limb bones, killing chondrocytes and mostly sparing the right limbs (Ahmadzadeh et al., 2020). While continuous expression of DTA from E12.5 generates extreme asymmetries by birth (not shown), transient activation from E12.5 to E17.5 (see Methods) leads to a subtler asymmetry (Fig. 6A), well suited to test the sensitivity of BASILISC. We scanned eight P3 mouse pups (four control and four experimental ones) and applied the Mimics version of BASILISC to measure left and right humerus, ulna, radius, femur and tibia. We then calculated the left/right ratio for each bone, and compared this parameter for all bones between experimental and control mice, via 2-way ANOVA (Fig. 6B). This analysis revealed a significant effect of the Genotype on limb asymmetry, with asymmetries ranging from ~5 to 20%, similar to our previous report (Ahmadzadeh et al., 2020). These results suggest that BASILISC can be used to quickly detect and characterize skeletal phenotypes affecting the long bones.

**Figure 6.**
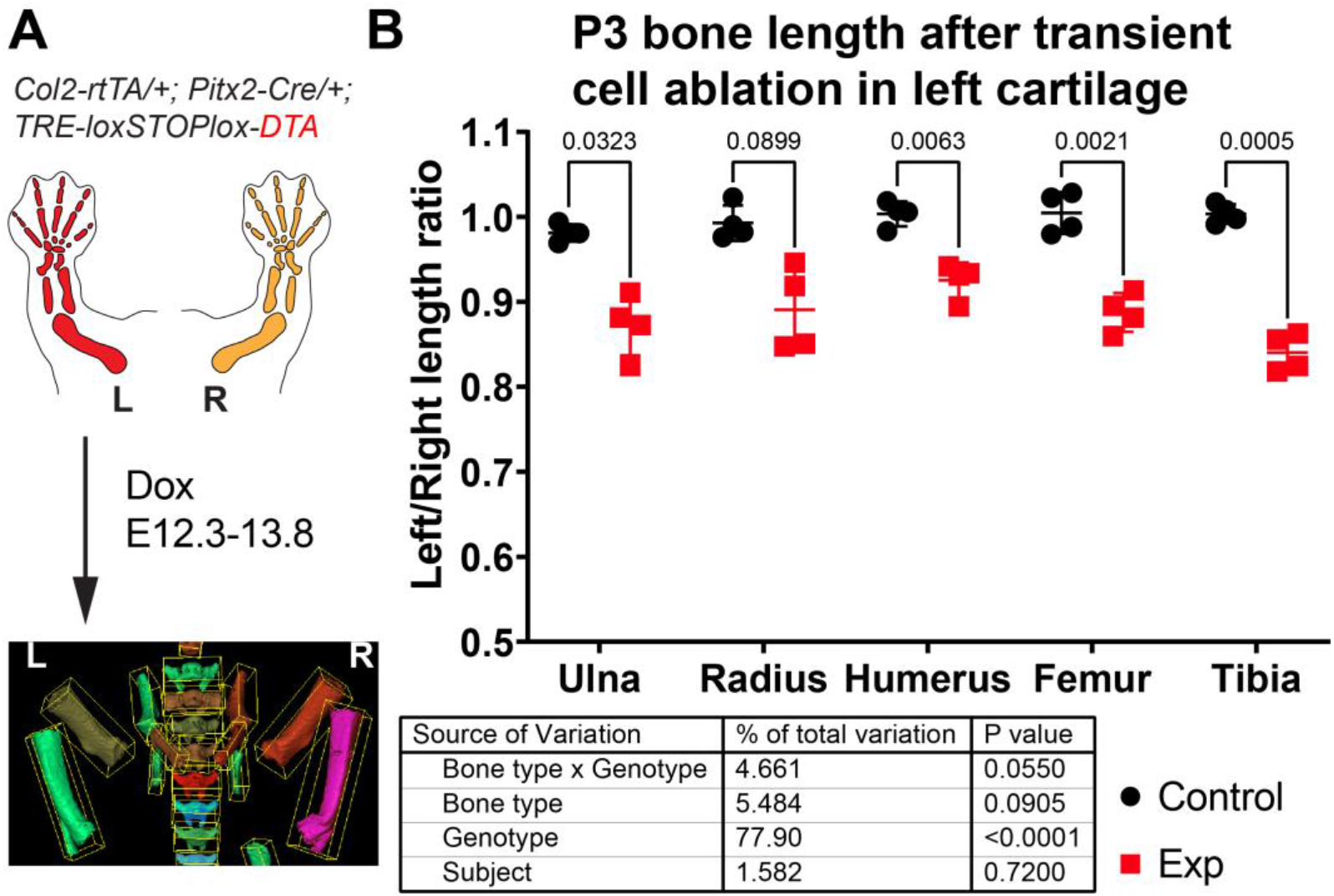
Application of BASILISC to a mouse model of limb asymmetry. A) Mouse model of transient cell ablation in the left cartilage. L, R: Left, Right. B) Left/right length ratios of the shown bones at P3. The table shows the results of repeated measures 2-way ANOVA (i.e. multiple bones analysed per sample), and the numbers in the graph are p-values for Sidak’s multiple comparisons post-hoc test.

## DISCUSSION

Here we have presented a fast and easy method to determine calcified bone length from XMT scans of whole mouse samples, without the need for dissecting the limbs, skinning or eviscerating the bodies. We tested our algorithm on a range of developmental stages (E17.5 through P7) that covers 9 days of very fast growth (Sanger et al., 2011).

### Advantages over classic skeletal preparations

As any developmental biologist working on limb patterning and/or growth has experienced, analysing one litter’s worth of samples by the classic method of skeletal preparation, limb microdissection, photograph acquisition and length measurement on the 2D pictures takes at least ten days and close to twenty hours of dedicated hands-on work (Rigueur and Lyons, 2014). With the BASILISC approach, decapitation and storage of the mouse bodies takes just a few minutes per litter; scan time is roughly ten minutes per sample (plus thirty minutes of set up per imaging session); data loading and analysis takes ~5 minutes per scan. On average, this amounts to 3-4 hours of hands-on work per litter. Another advantage is that the measurement is three-dimensional, as opposed to two-dimensional, and therefore robust to orientation errors. Lastly, the scan is not destructive, and therefore the samples can be later on processed for histology or other procedures (Baier et al., 2019; Hopkins et al., 2015).

### Comparison to previous automation approaches

In principle, the ideal pipeline for the kind of analysis that we perform here would be a fully automated method that recognised each of the long bones from a full-body scan, measured their length and exported those measurements without user intervention. In fact, there have been very impressive attempts at achieving this goal, combining object-based image analysis (that utilises shape and context-dependent information in addition to pixel intensity values) with machine learning. For example, Heidrich et al. (Heidrich et al., 2013) used Cognition Network Technology to extract objects and their properties from XMT data of chicken embryos at multiple stages, and then used these data to train a machine learning tool for automatic long bone classification. BASILISC is obviously far from achieving that level of automation, and while ImageJ is progressively including more machine learning modules, Mimics does not currently allow it. However, one of the strengths of BASILISC stems from its simplicity, as it can be used directly on any set of data, with minimal modification of the script. Contrary to this simplicity, the pipeline described in (Heidrich et al., 2013) required a large training set of close to 3,000 instances, and also complex iterative thresholding methods. Moreover, although the classification achieved via this complex process was remarkably accurate, it still required supervision and was only applied to a reduced developmental window.

In contrast to other automation procedures where edge detection has been used to determine optimal thresholds (Rathnayaka et al., 2011; Zhang et al., 2010), here we rely on a global threshold optimized by trial and error to find an optimal range of grey values. BASILISC could be further refined by implementation of widely used edge detection algorithms to further improve the segmentation process and potentially increase the accuracy of the measurements obtained. However, since intensity can vary across the length of long bones (Rathnayaka et al., 2011), edge detection would require the use of multiple thresholds to reduce the degree of error in segmentation. Thus, here we opt for a single global threshold to extend the capabilities of the algorithm for a range of developmental stages.

To our knowledge, our Mimics script is the first algorithm that makes use of the Python library within the Mimics software to automate the segmentation, 3D modelling and analysis of length of skeletal elements. Previously, Mimics has been complemented with other scripting languages like MATLAB (Huhdanpaa et al., 2011) for image processing before segmenting the data, or software like Creo elements (Rios et al., 2014) to analyse scans after they have been segmented. In the latter case, though, the reference points for length measurement had to be manually selected, which is a time-consuming step to do in 3D. Through BASILISC, segmentation and length measurements can all be obtained within the one program (be it Mimics or ImageJ) and extensive programming knowledge is not required. Furthermore, we provide a processing pipeline that goes from optimized scanning conditions of mouse samples across a range of developmental stages, to streamlined image processing and data analysis, making BASILISC a readily available tool for the research community.

It should be noted, however, that the BoneJ version of BASILISC is less accurate than the Mimics one (Fig. 2 and 3), and this could lead to some limitations in its applications. We hypothesise that BoneJ is more sensitive to discretisation and thresholding artefacts because it uses binary pixel values for input, while Mimics makes a mesh over greyscale data, smoothing out random bits of noise at the crucial areas on the bones’ extremities.

### Comparison with ‘real’ length measurements

Strictly speaking, the ‘true value’ of bone length cannot be obtained with absolute certainty by any method, as no measurement is devoid of error. For example, the classic skeletal preparation method involves quite a harsh procedure, including increasing gradients of glycerol that can shrink the sample up to 3-6% (Mabee et al., 1998). However, given the widespread use of skeletal preparation, flat mounting and imaging to obtain 2D length estimations, we compared the measurements obtained by the BASILISC approach (Mimics version) with the length obtained by skeletal preparations (Rigueur and Lyons, 2014). Of note, all conditions showed remarkable correlation between both methods, with 20-μm resolution and 398-HU threshold yielding measurements very well correlated to those obtained via skeletal preparations across the whole range of lengths analysed. One important consideration of the Mimics version is that the ‘line to surface’ fitting method generates the longest possible distance, which in some cases is not strictly running parallel to the element’s main axis (e.g. Fig. 2F). This obviously generates a small bias in the measurement, but as long as the same method is used to compare different experimental conditions, this bias will be consistent and is not expected to contribute to the observed biological effect. Similarly, the aligned box approximation used in our BoneJ version of BASILISC can also potentially lead to slight length overestimation, depending on the shape of the bone. Importantly, we also showed that the optimal Mimics and BoneJ versions of BASILISC yield highly correlated measurements when applied to the same scans.

### Limitations and future improvements

Some long bones are often segmented together in our pipeline, most often tibia and fibula (Fig. 2F and 3F), and sometimes radius and ulna. This is because their automatic separation would require too high a threshold. While radius and ulna can often be quickly separated manually using the *Split_mask* function in Mimics (see Methods), this is not feasible for the tibia and fibula, because their interaction surface is too large. This issue has some impact on the tibial measurements, because the fibula protrudes a bit farther than the tibia on the distal end (Fig. 2F and 3F). However, the effect is quite minor and we showed that under the right conditions the error is very consistent, as the left/right ratio for the tibia is quite tightly centred on 1 (Fig. 4). Therefore, the slightly overestimated tibial lengths can still be used for comparison purposes between different genotypes and/or treatments. The decision to invest more time in splitting them as opposed to accept the error is up to the user and depends on two main aspects: the degree of accuracy desired and the time investment required to correct the error in all samples. In our case, we opted not to correct this segmentation error, as the minor gain in accuracy would be outweighed by the extra time investment.

The aforementioned limitation would be corrected with an automatic classification system based on machine learning (Heidrich et al., 2013), but the implementation of these methods is not straightforward. If this capability is implemented in the future, it could speed up image processing even further, as in theory no user intervention would be required to seed landmarks and/or label the bones of interest.

### Other applications

The current BASILISC pipeline in Mimics only measures length of the elements, because it applies the ‘fit line to surface’ tool to the 3D models of the bones. However, it could in principle be adapted to measure width, by fitting a cylinder to the model and interrogating the width of the cylinder. This approach would require careful selection of the fitting parameters, so that the surface of the cylinder coincides with the surface of the 3D model. The BoneJ version, in turn, can directly be used to measure bone width, just by using the other dimensions of the aligned box.

## SUPPLEMENTARY INFORMATION

**Supplementary Video 1 caption.** Overview of the BASILISC process in Mimics, performed on one of the scans used for this study. The messages prompted by the script were taken as screenshots and added to the video clip. Please note that an artefact of the video capture causes the mouse cursor to be shown slightly displaced from its real position.

**Supplementary Figure 1. A)** Representative 3D viewer result window after applying the *Particle Analyser* tool to a segmented and cleaned up whole-body scan (P7 mouse). **B-B”)** Close-ups of femur (B), tibia (B’, right) and humerus (B’, left and B’’) showing the features that can be obtained from the *Particle Analyser* tool, as indicated. Green arrowheads point to the ends of the maximum Feret’s diameter. The red arrowhead points to a region where the longest principal axis does not align with the skeletal element. **C)** Two different samples were scanned four times at 40-μm resolution, and each of those scans were analysed twice, with identical or nearly identical parameters. **D)** Two different samples were scanned four times at 20-μm resolution, and most of those scans were analysed once. **E)** Heatmap with the average coefficient of variability for each bone and resolution.

**Supplementary Figure 2. Coefficients of variability for P7 and E17.5 analysis using Mimics and BoneJ.** A, B) Heatmaps for the CVs of the Left/Right length ratios obtained after analysis of P7 bones with Mimics (A) and BoneJ (B) pipelines. C, D) Heatmaps for the CVs of the indicated ratios obtained after analysis of E17.5 bones with Mimics (C) and BoneJ (D) pipelines.

## MATERIALS AND METHODS

### Animal experiments

Mouse embryo and pup samples were scavenged from other experiments in the Rosello-Diez lab, approved by the Animal Ethics Committee at Monash University (protocol 17048). Wild-type E17.5 samples were obtained from Asmu:Swiss crosses. P7 samples consisted of tTA-negative littermates (phenotypically wildtype) obtained from crosses of females containing the left-lateral plate mesoderm specific *Pitx2-Cre* (Shiratori et al., 2006) and a cartilage-specific *Col2a1-tTA* (Rosello-Diez et al., 2018) with males bearing a *Tigre^Dragon-DTA^* allele (Ahmadzadeh et al., 2020). P3 samples consisted of pups obtained from crosses of females containing the left-lateral plate mesoderm specific *Pitx2-Cre*, a cartilage-specific *Col2a1-rtTA* (Posey et al., 2009) and 1 copy of an *Egr1* null allele (JAX#012924) with males bearing a *Tigre^Dragon-DTA^* allele (Ahmadzadeh et al., 2020). Control and experimental animals were separated based on rtTA genotype, regardless of the presence of the *Egr1* null allele. Doxycycline hyclate (Sigma, 0.5 mg/ml in the drinking water, with 0.5% sucrose to increase palatability) was given to the pregnant female from E12.3 to E13.8. Noon of the day the vaginal plug was detected was considered E0.5.

### Micro-CT scans

A Siemens Inveon PET-SPECT-CT Small Animal scanner in CT modality was used for all experiments. Parameters: 20- and 40-μm resolution, 360 projections at 80 kV, 500 μA, 600 ms exposure with a 500ms settling time between projections. Binning was applied to vary resolution with 2×2 for 20μm and 4×4 for 40μm scans and data was reconstructed using a Feldkamp algorithm. The samples (beheaded embryo and pup bodies) were placed in supine position over custom-fitted foam bedding, so that the limbs were not in contact with any hard surface.

### BoneJ software and pipeline

FIJI was used to develop the analysis pipeline. See Results for an overview; each step in the process is outlined in detail below.

#### Data loading

The entire image set is loaded into memory, as many of the downstream actions requires to have the entire image set in memory. A virtual stack is possible as the initial action, if it is necessary to crop or reduce the size of the image, if the image is too large for the computer being used. When the crop/size reduction is performed, FIJI will load the entire object into memory anyways.

#### Segmentation

This method employs the segmentation method of maximum entropy. Segmentation divides the image into multiple parts (at least two, usually). This yields the objects of interest, and everything else. Some methods, such as Otsu’s three-class, can produce more than two layers. The choice of maximum entropy in this case is based on making the method as widely applicable as possible. Maximum entropy is implemented in ImageJ as a method to maximise the inter-class entropy (between the selected objects and everything else in the image). This involves the average grey values of the pixels present in each image (or in a reference image), the individual grey value of each pixel, and the grey values in the local neighbourhood of each pixel. The initial threshold value the algorithm selects is based on the probability estimation from a histogram of all pixels in the image stack. Therefore, maximum entropy offers the best robustness, rejection of undesirable elements, acceptable performance in low-contrast images, and good noise tolerance. The method is described in (Kapur et al., 1985). Threshold values of 280 to 500 HU were chosen, depending on the developmental stage, as a best compromise on the images included. This value may need to be adjusted depending on the CT scanner, reconstruction kernel, and the sample. The segmentation was further refined via the *Erode* and *Watershed* tools (2D versions).

#### Generation and labelling of aligned boxes with *Particle Analyser*

BoneJ’s *Particle Analyser* code was altered to use the bone’s inertia tensor to define the three principal axes of each element, and then generate the minimum-size cuboidal box that fits the skeletal element and that is aligned with the principal axes. The coordinates and dimensions of these boxes are then appended to the Results window in *Fiji*. The *Label Elements* (*3D*) feature then allows the user to Ctl-click the surfaces of interest and assign a label to them. These are added as a new column to the *Results* window. The macro below also includes the possibility of saving a screenshot of the *3D Viewer* window with the desired orientation.

#### ImageJ macro to automatize the process

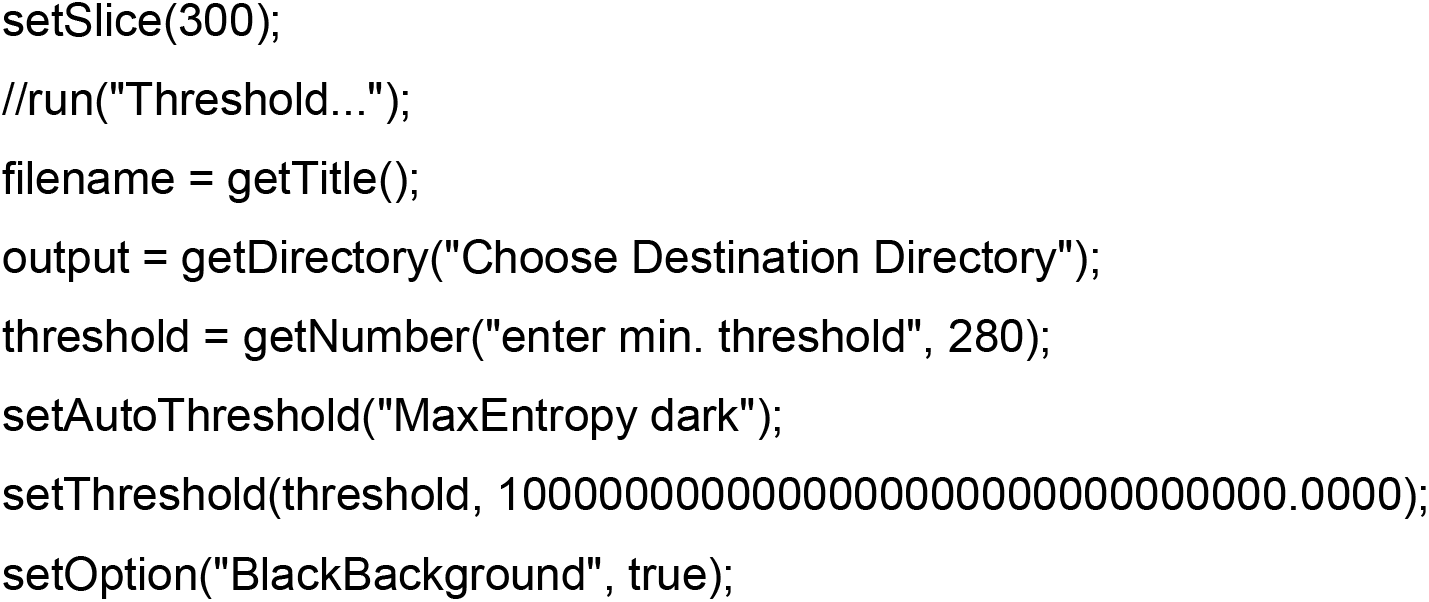
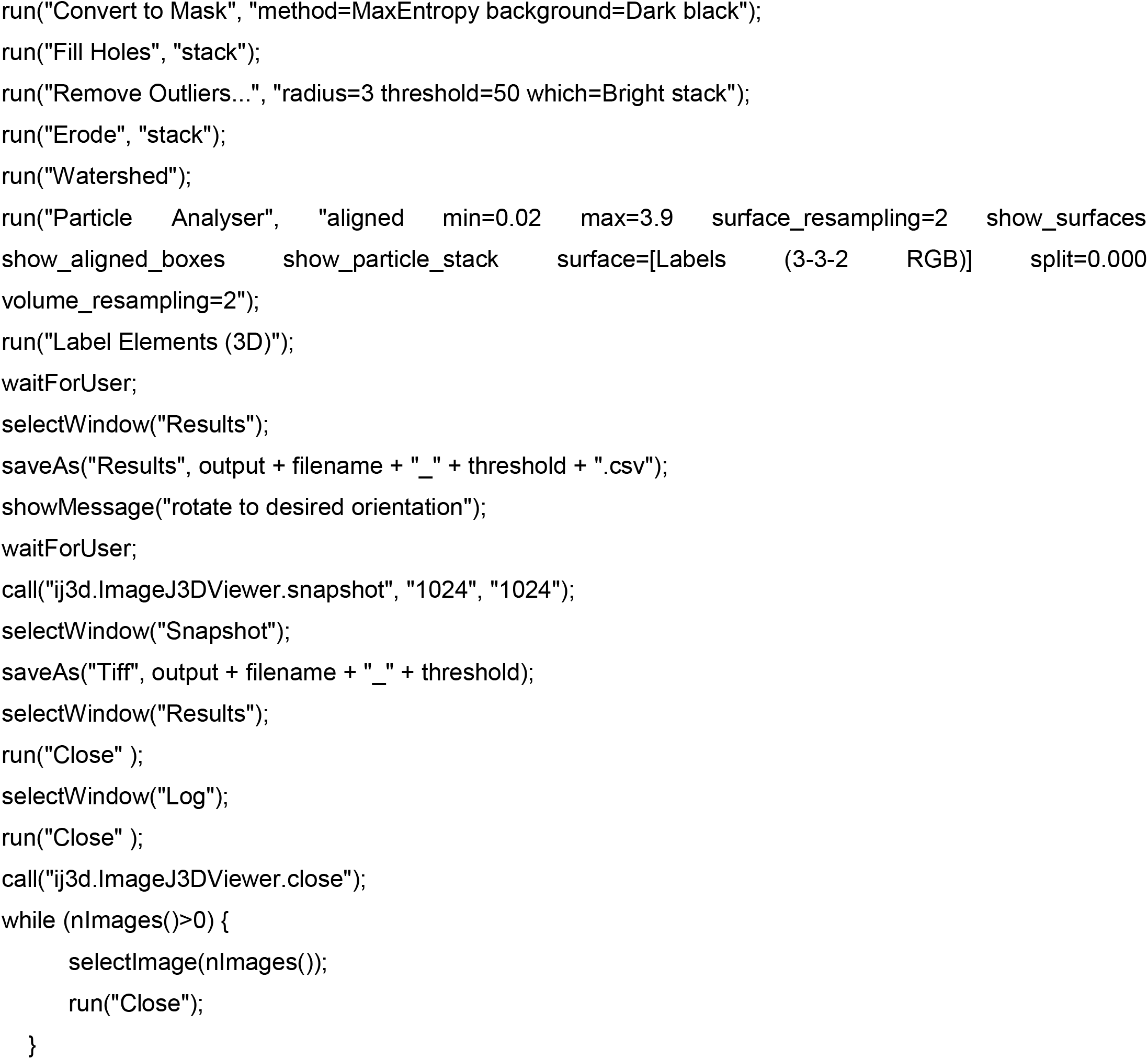

#### R-script to merge and clean-up data tables

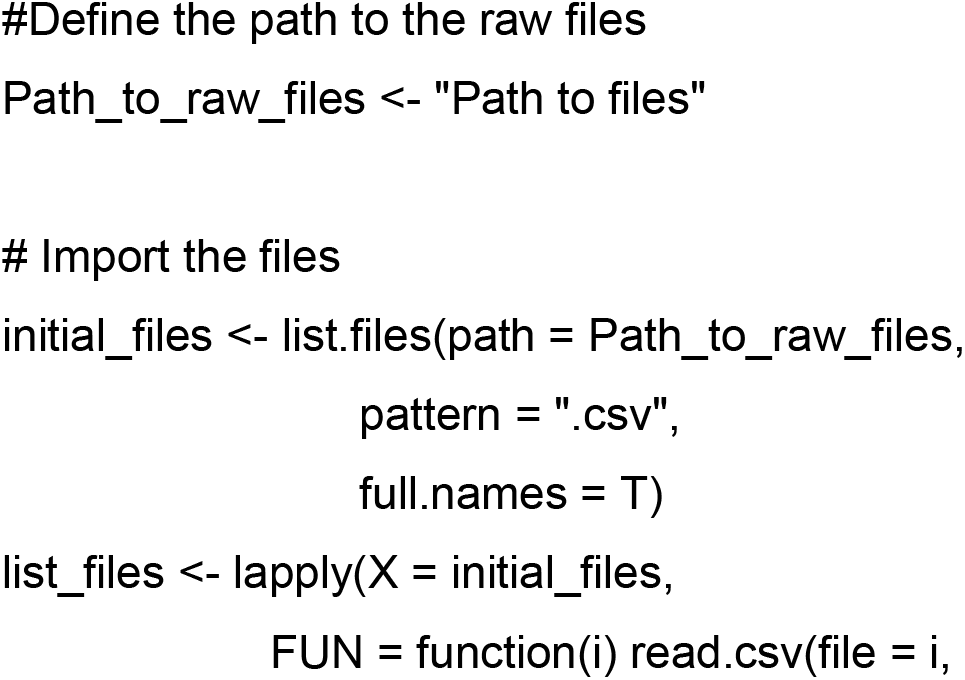
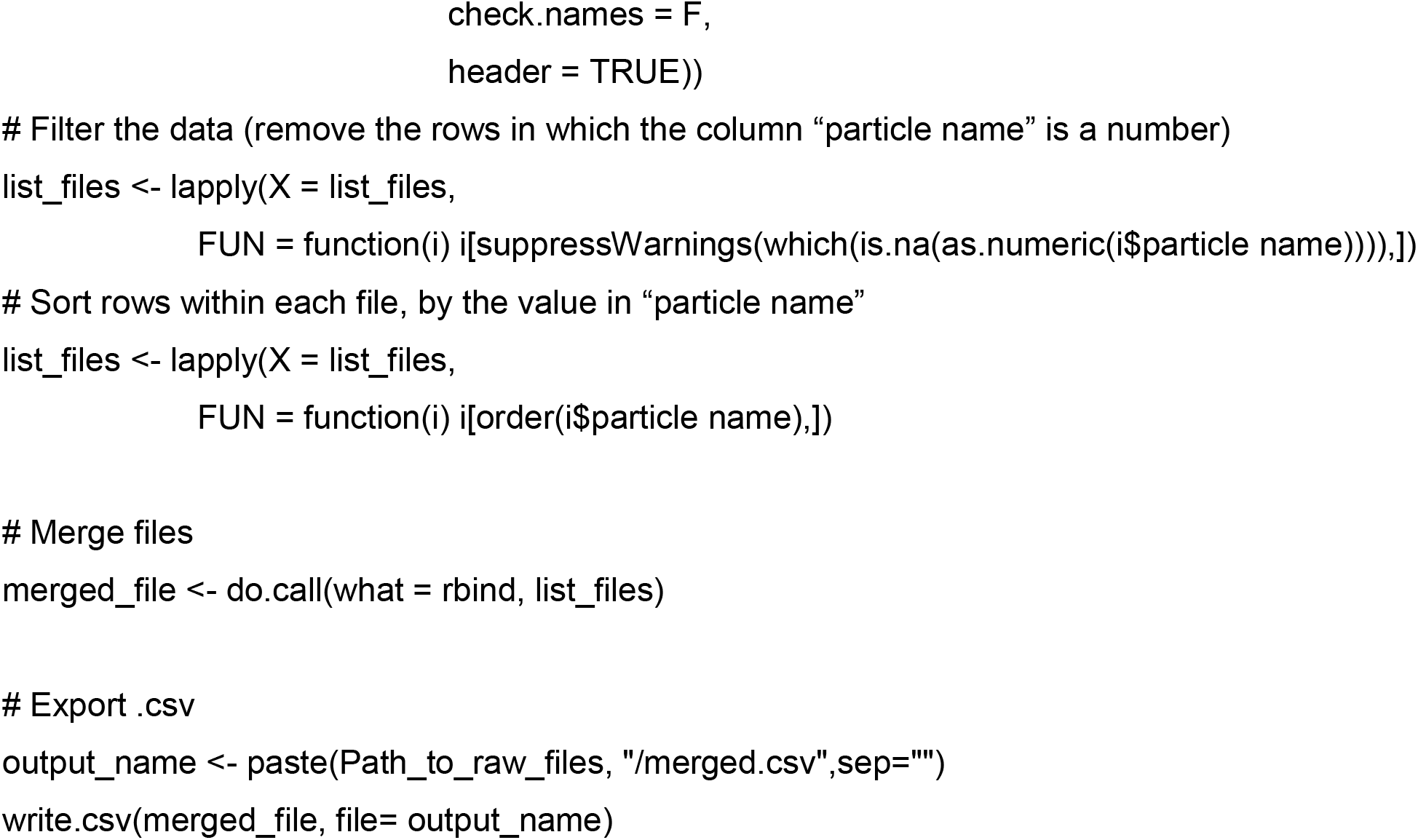

### Mimics software and pipeline

Mimics Research (v21.0; Materialise, Leuven, Belgium) equipped with the scripting module was used to develop the analysis pipeline and the Python script described here. See Results for an overview and each step in the process is outlined in detail below.

#### Thresholding

As soon as the DICOM data is uploaded into Mimics, the first step is to distinguish the bones from all the other tissue by defining a range of Hounsfield Units (HU) that corresponds to bone density. The first Python command in BASILISC creates a mask labelled “ALL”, which will segment all the skeletal elements present, creating a global threshold specific to this mask. Since the goal was to measure the developing mineralised part from end to end, this step had to detect immature trabecular bone at the ends of the growing elements. In our uncalibrated XMT scans, we realised that the custom minimum threshold for bone tissue defined by Mimics (226 HU) often over-represented the actual bone tissue in the scans as it selected a greater area of tissue. The optimal lower threshold for the developmental stages of interest had thus to be determined empirically. Although Mimics can take both gray scale values (GV) and HU units, the input in the script can only be into GV, and therefore the first step was to transform the data into GV to adequately segment all bones from the rest of the tissue. This is achieved through the “segment” attribute seen on the last line of code for this section. In principle, different optimal thresholds exist for different scanning conditions and certainly for different developmental stages, as the ratio between woven and lamellar bone decreases, and hence BASILISC was designed in such a way that the user can select among three pre-defined thresholds via a pop-up menu. This can be easily changed within the following section of the script (pre-defined values appear in orange font):

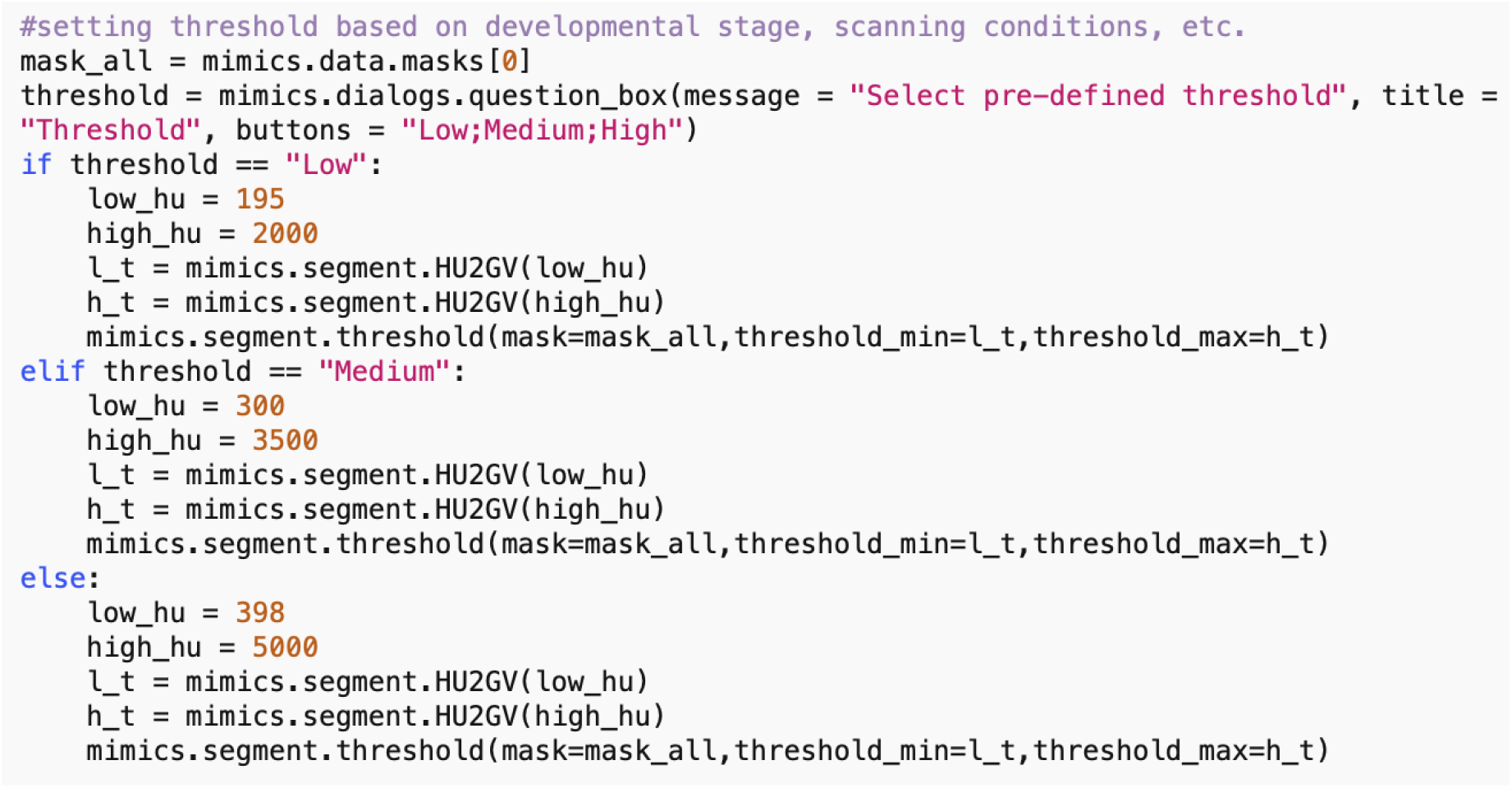

#### Landmarking

The purpose of this step is to segment and uniquely label all the bones of interest, using Mimics tools. This is achieved using a function that prompts the user (via a pop-up window) to select a landmark on the bone of interest. The first step in landmarking is to select the bones of interest to create a list of “landmarks”. This list contains the unique name of each selected bone and defines the order of segmentation during the process. The “indicate_landmark” function guides the user through each of the bones to be segmented by means of a dialogue box, asking the user whether a given element is present in the scan or not and with two active buttons: “Select” & “Skip” (Fig. 1C). The user has the option to skip an element if a given bone is not present in the scan, this would then be excluded from the analysis. If the “Select” button is activated, a second dialogue box prompts the user to select a region (landmark) of the indicated element by simply clicking on it on one of the 2D views of the sample. BASILISC will automatically label and segment the selected element without further user interaction, through the Mimics “region_grow” function. A FOR loop has been included in the BASILISC script when executing the “indicate_landmark” function, so that the steps above are recursively followed <u>for</u> each of the bones of interest sequentially, using the name of each bone as an index within the FOR loop.

#### Generation of 3D models

Once all the elements of interest have been segmented and labelled accordingly, a function has been created in BASILISC that creates 3D models of each element, “create_3D”. A FOR loop in the script steps through each of the segmented bones and creates a 3D model of each at the highest possible resolution (Fig. 1D). This provided the most accurate measurements possible and since a limited number of bones are analysed, computing time to create each 3D model did not increase significantly.

#### Measurement

Once BASILISC has automatically made 3D models, it will fit a centre line to each bone within the “create_3D” function. This is achieved through the “analyze.create_line_fit_to_surface” attribute in Mimics. The script has been designed to then automatically obtain the length of the fitted line and save the measurement in a text file (Fig. 1E). Since this step is included within the function described above, which includes a FOR loop, the line is fitted as each 3D element is made, and the measurement is recorded progressively. The text file created will have the name of the given part, e.g. RIGHT HUMERUS, followed by a comma and the corresponding length of the element. This step is done automatically without any user input required after the landmarking step has been finalized. As the file created is only labelled with the name of the developmental stage created, the user should change the name of the text file to be sample specific before analysing the next sample.

### Manual corrections during image analysis

For E17.5 samples, the radius and ulna are segmented together at the thresholds we use, but they could be easily separated using the *Split mask* function of Mimics, as their interaction surface was quite reduced. This required identification of specific areas where there was an overlap in pixels, manually selecting areas that corresponded to each bone in a 2D view in at least three areas across the length of the bone (proximal, distance and middle area), before extrapolating the selected regions to create two separate elements. This tool is not available in BoneJ.

### Pipeline benchmarking

For Figures 2 and 3, each specimen was scanned in triplicate or quadruplicate (on three or four different days), at two resolutions each (20 and 40 μm), and each of the 6 scans was segmented at two different lower thresholds (in Mimics: 650 and 398 HU for P7, 398 and 226 HU for E17.5; in BoneJ: 500 and 385 HU for P7, 400 and 280 HU for E17.5) to perform length measurements. Humerus, radius, femur, tibia and clavicle (the latter only for E17.5) were analysed for two (P7) or three (E17.5) different specimens.

### Skeletal preparations

After embryo collection, the skin, internal organs and adipose tissue were removed. The samples were then fixed in 95 % EtOH overnight at room temperature. To remove excess fat, the samples were then incubated in acetone overnight at room temperature. To stain the cartilage, the samples were submerged in a glass scintillation vial containing Alcian blue solution (0.04 % (w/v), 70 % EtOH, 20 % acetic acid) and incubated at least overnight at room temperature. The samples were destained by incubating them in 95% EtOH overnight, and then equilibrated in 70% EtOH, prior to being pre-cleared in 1% KOH solution for 1-10h at room temperature (until blue skeletal elements were seen through). The KOH solution was replaced with Alizarin red solution (0.005 % (w/v) in 1% KOH) for 3–4h at room temperature. The Alizarin red solution was then replaced with 1-2% KOH until most soft tissues were cleared. For final clearing, the samples were progressively equilibrated through 20% glycerol:80% (1%KOH), then 50% glycerol:50% (1% KOH) and finally transferred to100% glycerol for long-term storage.

## Acknowledgements

We thank Jonathan Bensley for initial efforts with the initial BoneJ protocol, Adrian Salavaty for help with the generation of the R script, Hyab Mehari Abraha (Panagiotopoulou lab) for Mimics training and Noramira Azlan (Materialise) for providing a Mimics trial license for the initial steps of the project. The authors acknowledge the facilities and scientific and technical assistance of the National Imaging Facility (NIF), a National Collaborative Research Infrastructure Strategy (NCRIS) capability at Monash Biomedical Imaging (MBI), a Technology Research Platform at Monash University. We acknowledge the technical assistance of Tara Sepehrizadeh who is a NIF Facility Fellow.

