## Supplementary figures and images for "A new pipeline to automatically segment and semi-automatically measure bone length on 3D models obtained by Computed Tomography"

### Suppl. Fig. 1

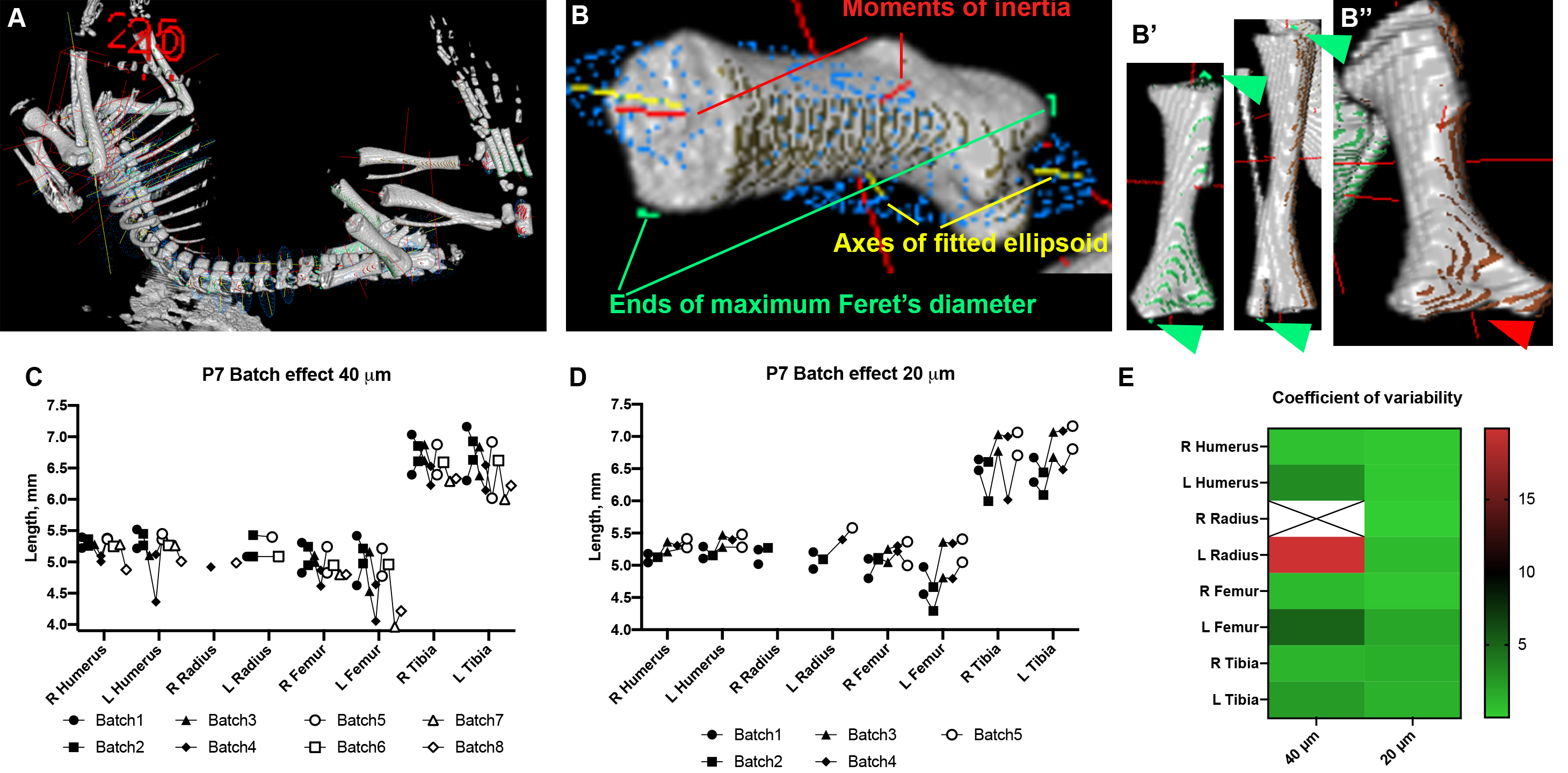

### Suppl. Fig. 2

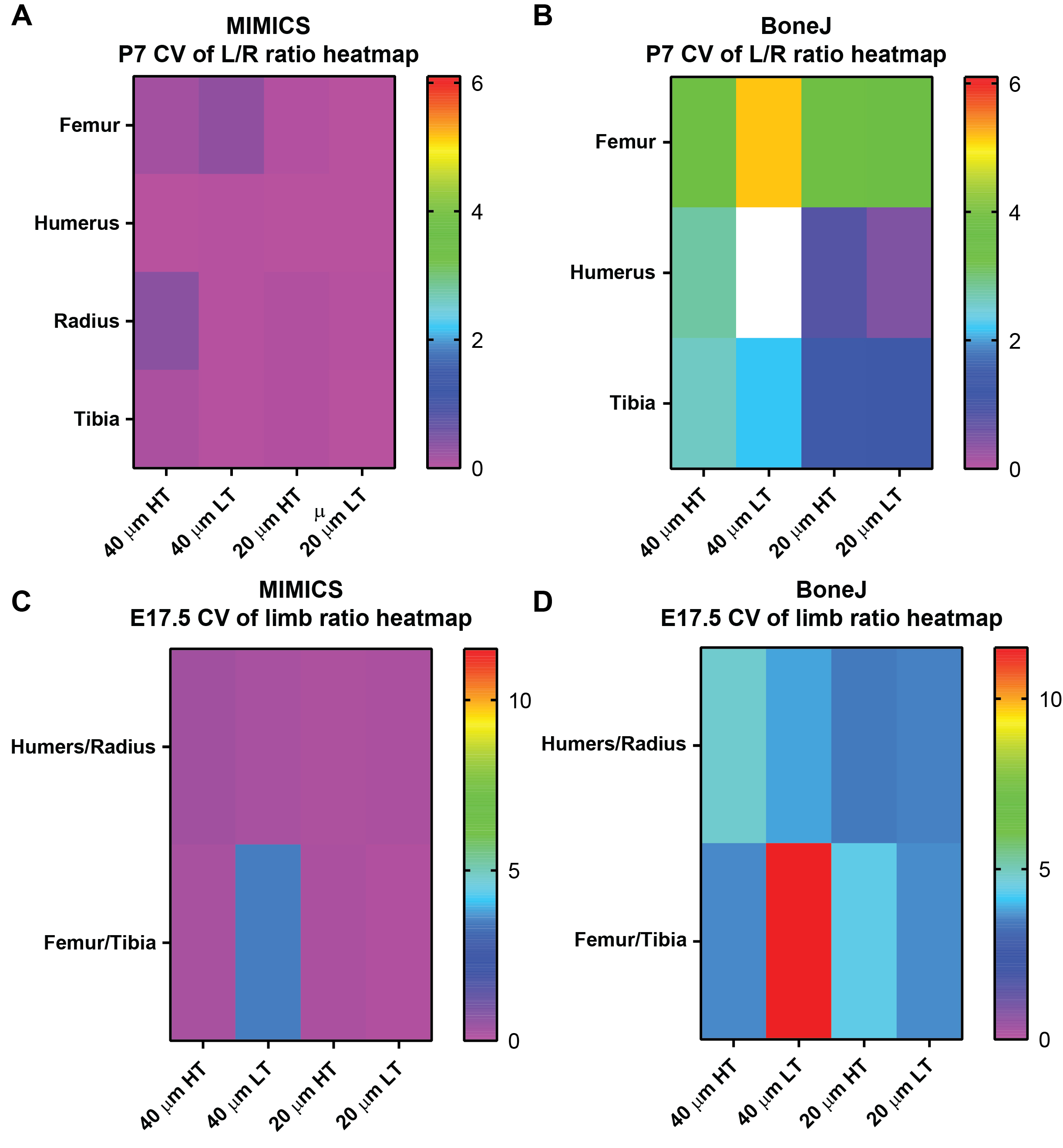
